# Lipid rafts interaction of the ARID3A transcription factor with EZRIN and G-Actin regulates B cell receptor signaling

**DOI:** 10.1101/2020.11.11.378489

**Authors:** Christian Schmidt, Laura Christian, Josephine Tidwell, Dongkyoon Kim, Haley O Tucker

## Abstract

Previously we demonstrated that the ARID3A transcription factor shuttles between the nucleus and the plasma membrane, where it localizes within lipid rafts. There it interacts with components of the B Cell Receptor (BCR) to reduce its ability to transmit downstream signaling. We demonstrate in this report that a direct component of ARID3A-regulated BCR signal strength is cortical Actin. ARID3A interacts with Actin exclusively within lipid rafts via the Actin binding protein EZRIN, which confines unstimulated BCRs within lipid rafts. BCR ligation discharges the ARID3A-EZRIN complex from lipid rafts allowing the BCR to initiate downstream signaling events. The ARID3A-EZRIN interaction occurs almost exclusively within unpolymerized G-Actin where EZRIN interacts with the multifunctional ARID3A REKLES domain. These observations provide a novel mechanism by which a transcription factor directly regulates BCR signaling.

## INTRODUCTION

The cortical cytoskeleton is an Actin-based structure that underlies and confers mechanical support to the plasma membrane (PM) via a dense meshwork (∼100 nm thick) comprised primarily of filamentous (F) Actin (1). An array of proteins, including myosin motors as well as Actin-binding and linker factors comprise the PM (2). The cortex is dynamic in that it can locally disassemble (G-Actin) and reassemble (F-Actin) to drive formation of cellular protrusions and to modulate lipid microdomain (lipid raft) organization (3, 4). PM-associated receptors maintain dynamic contacts with the cortex via linker proteins that simultaneously bind Actin and the cytoplasmic domains of PM proteins (5). The resulting receptor-cytoskeleton anchoring has a direct impact on receptor mobility and influences receptor function (6). Over 100 Actin-binding proteins constitute the cortical cytoskeleton (1).

In resting B cells, the mobility of B Cell Antigen Receptors (BCRs) is restricted, as the cortical Actin cytoskeleton acts as a barrier to their diffusion (7, 8). Such contractile stress and tension is mediated by ERM proteins (<u>E</u>ZRIN, <u>R</u>ADIXIN and <u>M</u>OESIN) (9, 10). ERM proteins supply the tension needed to anchor B Cell Receptors (BCRs) in the cortex. Actin and EZRIN form a network which confines BCRs in nanoscale lipid rafts via EZRIN carboxyl terminal binding to integral membrane proteins of the BCR complex (11). BCR mobility, on the other hand, is established if the EZRIN-Actin interaction is disrupted (12).

Besides its mechanical functions, Actin has also been implicated in transcriptional regulation— both through cytoplasmic alterations in cytoskeletal ACTIN dynamics and through the assembly of transcription factor (TF) regulatory complexes (13). Established examples include the subcellular localization of myocardin-related TFs (MAL, MKL1, BSAC and MRTF-B) (14, 15), the PREP2 homeoprotein TF, and the TF repressor, YY1 (16, 17).

We demonstrated (18) that a palmitoylated pool of ARID3A, a B-cell/embryonic stem cellrestricted TF, shuttles from the nucleus to the PM where it is diverted to lipid rafts of resting B cells to associate with “signalosome” components. The BCR signalosome requires participation of signaling proteins (eg, BTK and BLNK) whose genetic defects often impair B cell activation, differentiation and often lead to agammaglobulinemia (11, 12). Following BCR stimulation, ARID3A transiently interacts with SUMOylation enzymes, blocks calcium flux and phosphorylation of BTK and TFII-I TFs and is then discharged from lipid rafts as a SUMO-I-modified form (18). The lipid raft concentration of ARID3A contributes to the signaling threshold of B cells, as their sensitivity to BCR stimulation decreases as the levels of ARID3A increase (18). These events are independent of the previously established role of nuclear ARID3A in immunoglobulin gene transcription (19, 20).

We concluded from the results above that ARID3A contributes to a BCR tuning mechanism within lipid rafts, yet further details of how this mechanism is initiated at the PM remained unknown. Here we demonstrate that a critical component underlying ARID3A-regulated BCR signal strength is Actin. ARID3A interacts with Actin, exclusively within lipid rafts, via the Actin-binding protein, EZRIN, which in unstimulated BCRs is confined within lipid rafts. We observed that antigen binding discharges the ARID3A-EZRIN complex from lipid rafts allowing initiation of downstream signaling events. Consistent with the timing of EZRIN discharge, ARID3A-EZRIN interaction occurs almost exclusively within unpolymerized, G-Actin, and is mediated by the multifunctional REKLES domain of ARID3A. These along with our previous data provide further mechanistic insight as to how TFs can directly regulate BCR signaling strength.

## RESULTS and DISCUSSION

### Variable levels of ARID3A within lipid raft in unstimulated and stimulated B cell lymphomas

From B cell leukemias/lymphomas, we prepared whole cell lysates (WCL) as previously described (18). Lipid rafts were prepared by flotation on discontinuous sucrose gradients (18) using the B cell-specific lipid rafts component, Raflin (21), as an internal control for purity (Materials and Methods). Plasma membranes (PM), isolated as pellets from the same raft extraction, were further purified for 30 min on ice as described in Materials and Methods.

As shown in S-Figure 1, we estimated that raft-localized ARID3A accounted for <10% of total cellular ARID3A—a value similar to the previously estimated level of membrane (m) IgM in lipid rafts (22, 23). Lipid rafts from Raji and Daudi Burkett lymphomas contained 10-fold less ARID3A than those of the CL01 B cell leukemia or the Ramos Burkett lymphoma (S-Fig. 1). Equal loading was demonstrated by probing the blots with anti-Raftlin. No significant differences were observed in either WCLs or M fractions. The differences among these lines in raft-localized ARID3A made them excellent choices for our analyses of anti-BCR signal strength.

### BCR stimulation induces a rafts-restricted, transient interaction of ARID3A with EZRIN prior to their co-discharge

We previously demonstrated that Ramos and CL01 were less sensitive, whereas Raji and Daudi were more sensitive to BCR stimulation (18). We further showed that engagement of the BCR results in a significant and specific reduction of the small pool of lipid raftslocalized ARID3A, which is lost from lipid rafts, as BTK and other signalosome components accumulate there in proportion to BCR signaling strength (18).

That the cytoskeletal linker protein EZRIN undergoes a similar anti-IgM-mediated discharge from lipid rafts (11, 12) prompted us to examine ARID3A-EZRIN interaction under differential BCR signal strength conditions. We observed that EZRIN and ARID3A did not form an immunoprecipitable complex in WCL of any of the B cell lines, regardless if the cells were stimulated or not [Fig. 1A(a-c), lanes 5, 6, 11, 12, 17, 18, 23 and 24]. However, EZRIN and ARID3A did associate within lipid rafts, and their interaction required BCR stimulation (compare lanes 7 and 19 with lanes 8 and 20 in Fig. 1A(a, b).

**Figure 1.**
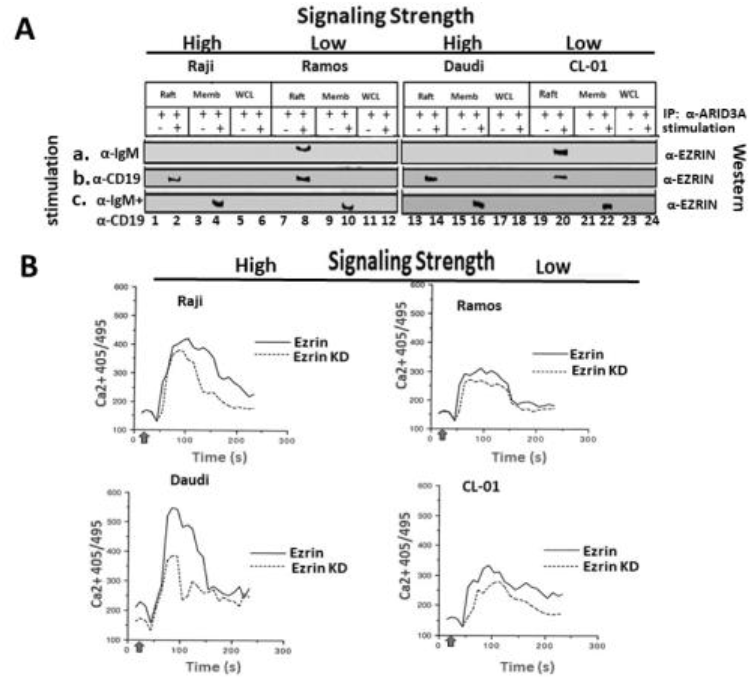
The ARID3A-EZRIN-complex is directed to different fates depending on the signaling threshold of the BCR. Raji, Ramos, CL01 and Daudi (10^8^ cells/dish) were stimulated for 5 min with (**a**)100 ng anti-IgM(α-IgM), (**b**) 100 ng α-CD19 or (**c**) 100 ng α-IgM +100 ng α-CD19. Lipid rafts (Raft), plasma membranes (Memb) and whole cell lysates (WCL) were analyzed following anti-ARID3A IP and Western blotting with anti-EZRIN. EZRIN-ARID3A do not Co-IP in either stimulated or nonstimulated WLC (lanes 5, 6, 11, 12, 17, 18, 23 and 24) but do within BCR-stimulated lipid rafts (<u>lanes 8 and 20</u> in **a** and **c**). ARID3A-EZRIN Co-IP in lipid rafts of the higher threshold cell lines (Ramos and CL01) stimulated with anti-IgM only (**a**, <u>lanes 8 and 20</u>); within all cell lines stimulated with anti-CD19 only (**b**, <u>lanes 2, 8, 14 and 20</u>); and are discharged in all cell lines following anti-IgM + anti-CD19 stimulation (**c**, <u>lanes 2, 8, 14 and 20</u>). EZRIN is retained in the Memb only following strong co-stimulation (**c**, <u>lanes 4, 10, 16 and 22</u>). **B**. EZRIN KD results in reduced signaling strength. EZRIN levels were reduced by shRNA knockdown (KD; S-Fig 4) and calcium (Ca2+) flux was measured in the indicated B cell tumors of high or low signaling strength following strong costimulation as detailed in Materials and Methods. Data shown are representative of 3 independent experiments. α-IgM + α-CD19 were added 30 s (upward arrow) before the beginning of the experiment, and the cells were analyzed at 250 cells/s. Results are plotted as the mean calcium concentration vs time. A significant difference was apparent between high vs low signaling cells when comparing their calcium response to stimulation with and without Ezrin knockdown.

EZRIN was co-discharged with ARID3A as evidenced by their identical trafficking. That is, EZRIN was detected in anti-ARID3A IPs of lipid rafts of the higher threshold cell lines (Ramos and CL-01) stimulated with anti-IgM only [Fig. 1A(a), lanes 8 and 20); within all cell lines stimulated with anti-CD19 only [Fig. 1A(b), lanes 2, 8, 14 and 20]; and discharged in all cell lines following anti-IgM + anti-CD19 [Fig. 1A(c), lanes 2, 8, 14 and 20] stimulation. Notably, EZRIN was retained in the membrane fraction only following strong co-stimulation [Fig. 1A(c), Lanes 4, 10, 16 and 22]. We observed no change in the expression levels of SUMO-1, which modifies ARID3A at a single residue and interacts with ARID3A in rafts (18; S-Fig. 2).

These results suggested that post-discharge, the ARID-EZRIN-complex is directed to different fates depending on the signaling threshold of the BCR.

### EZRIN knockdown results in reduced BCR signaling strength

The above results suggested that, within lipid rafts, EZRIN might link ARID3A indirectly to the BCR to control signaling strength. Thus, we tested if signal strength was impaired following (anti-IgM + anti-CD19) stimulation in B cells deficient in EZRIN.

Retroviral shRNA knockdown (KD) of EZRIN was assessed and validated in S-Figure 3 as 7590% relative to a scrambled sh-RNA KD control. EZRIN WT and KD B cell leukemias/lymphomas were loaded with Indo-1, incubated with an Fcγ III/II receptor-blocking antibody (to prevent antibody-antigen immune complexes), and then strongly stimulated with anti-CD19 + F(ab′)2-IgM 30 seconds (s) prior to measurement of Ca^2+^ flux. Cells were gated and representative results of three separate experiments are shown in Figure 1B. EZRIN loss led to significant reduction in signaling strength of the more sensitive Raji and Daudi B cells as compared to the less sensitive Ramos and CL01 B cells. That is, ∼38% of Raji and Daudi responded to strong BCR stimulation as compared to ∼12% of Ramos and CL01.

These data lead us to speculate that the ARID3A-EZRIN complex might function to secure the raft environment for the active BCR complex or to prepare inactive BCR complexes for signaling activity. They further implied that the inherent signaling strength of the BCR, in the absence of EZRIN, may contribute in a feedback manner by directing the ultimate fates of ARID3A-EZRIN complexes within the Actin cytoskeleton—a hypothesis tested below.

### ARID3A interacts via EZRIN with Actin

Previous reports established that EZRIN binds specifically to polymerized F-Actin (11, 12) and that partial depolymerization of Actin increased the strength of BCR signaling (24). This and the data above suggested that EZRIN mediates BCR ligation– dependent aggregation of lipid rafts by releasing them from the underlying cortical Actin cytoskeleton.

As shown in Figure 2A, Actin and ARID3A interact in lipid rafts of all unstimulated B cell lines (lanes 1, 5, 9 and 13). Strong stimulation with anti-IgM + anti CD19 (Lanes 4, 8, 12, and 16) discharged ARID3A from Actin in lipid rafts of all cells. Weaker stimulation with either anti-CD19 or anti-IgM discharged Actin from ARID3A in lipid rafts of strong signaling strength B cells (Raji and Daudi; Lanes 2, 3, 14 and 15), but not from weak signaling strength B cells (Ramos and CL-01; Lanes 6, 7, 10 and 11). Strong stimulation (Fig. 2A; lanes 4, 6, 12 and 16) was required to discharge from lipid rafts other ARID3A interacting proteins, including UBC-9, PIAS-1 and SUMO-I.

**Figure 2.**
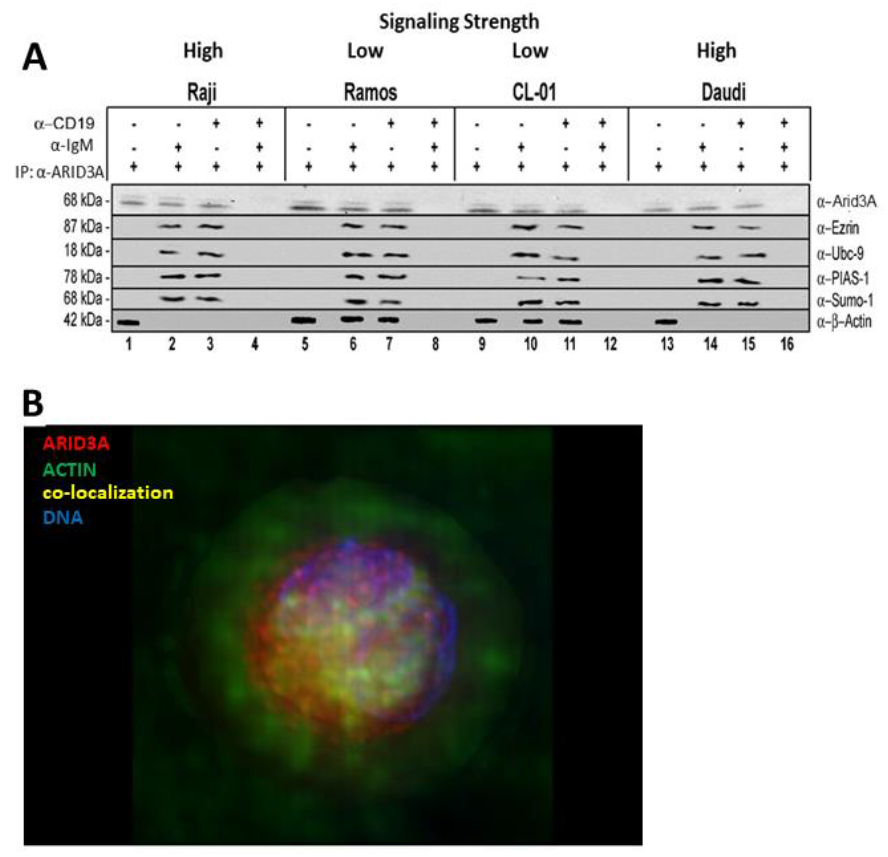
Interaction and discharge of ARID3A-Actin complexes within lipid rafts as a function of BCR signal strength. **A**. ARID3A Co-IPs with αβ-Actin in lipid rafts of B cell lines (<u>lanes 1, 5, 9 and 13</u>) regardless of their signaling strength. Strong stimulation (α-IgM + α-CD19; <u>lanes 4, 8, 12, and 16</u>) discharges ARID3A from ACTIN in lipid rafts of all cells, whereas weaker stimulation (α-CD19 or α-IgM) discharged ACTIN from ARID3A in lipid rafts of strong (Raji and Daudi; <u>lanes 2, 3, 14 and 15</u>), but not weak (Ramos and CL-01; <u>lanes 6, 7, 10 and 11</u>) signaling strength B cells. Strong stimulation (<u>lanes 4, 8, 12 and 16</u>) was required to discharge other ARID3A interacting proteins (UBC-9, PIAS-1 and SUMOI) from lipid rafts. **B**. Triple immunofluorescence of murine B cells stained for Actin (green), ARID3A (red) and nuclei (blue) revealed strong Actin-ARID3A association (yellow; also illustrated in the movies of SFigure 4)

That ARIA3A was detected in lipid rafts implied it was discharged after strong stimulation. However, our data do not exclude that other, unknown proteins also were discharged, as no antibodies against these putative factors were employed.

Support for an ARID3A-Actin interaction was provided by computerized 3D reconstruction of immunofluorescence data. As shown in Figure 2B, reconstructions (detailed in Materials and Methods) of strongly stimulated murine B cells stained for Actin (green), ARID3A (red) and nuclei (blue) revealed strong Actin-ARID3A association (yellow; also illustrated in the movies of S-Figure 4)

### ARID3A localizes within monomeric G-Actin

We reasoned that a functional consequence of ARID3A discharge is to release lipid rafts-associated Actin either for depolymerization or for polymerization. The degree to which release is achieved might contribute to the signaling threshold of the BCR.

In nonmuscle cells of mammals, Actin is encoded by two genes: *Actb* and *Actg1*, which encode beta-Actin and gamma-Actin respectively. Each isoform can exist as a monomer, termed Globular (G)-Actin, or as a polymer, Filamentous (F)-Actin (25, 26). Changes in Actin organization are driven by the assembly and disassembly of F-Actin. This dynamic turnover is regulated by a diverse set of proteins, many of which bind to F-Actin to influence the network architecture (27, 28).

Following transfection of ARID3A into COS7 cells, we employed Triton X-100 with or without TWEEN 20 fractionation to obtain detergent-soluble (enriched in G-Actin) and insoluble (enriched in FActin) proteins as described in Materials and Methods. Following lysate fractionation over SDS-PAGE, Western blotting was performed to determine the partitioning of ARID3A and EZRIN. Acid sphingomyelinase (SMPD1) served as a positive control for the insoluble (IS) fraction, and levels of input were adjusted using a pan-ACTIN Ab that reacts equally with both forms.

Unexpectedly, given that the vast majority of Actin-binding proteins associate with F-Actin (27, 28), ARID3A partitioned almost exclusively within the unpolymerized, G-Actin soluble fraction (Fig. 3A). These unexpected results encouraged us to analyze Actin-associated partitioning of a highly conserved, ARID3A paralogue, ARID3C (29). ARID3C is significantly condensed relative to ARID3A (S-Fig. 6), but both share all functional domains and undergo nuclear-cytoplasmic shuttling, with a fraction of ARID3C also localizing within lipid rafts following BCR stimulation (26). These shared features provided the opportunity to readily and more broadly investigate essential sequences required for G-Actin partitioning

**Figure 3.**
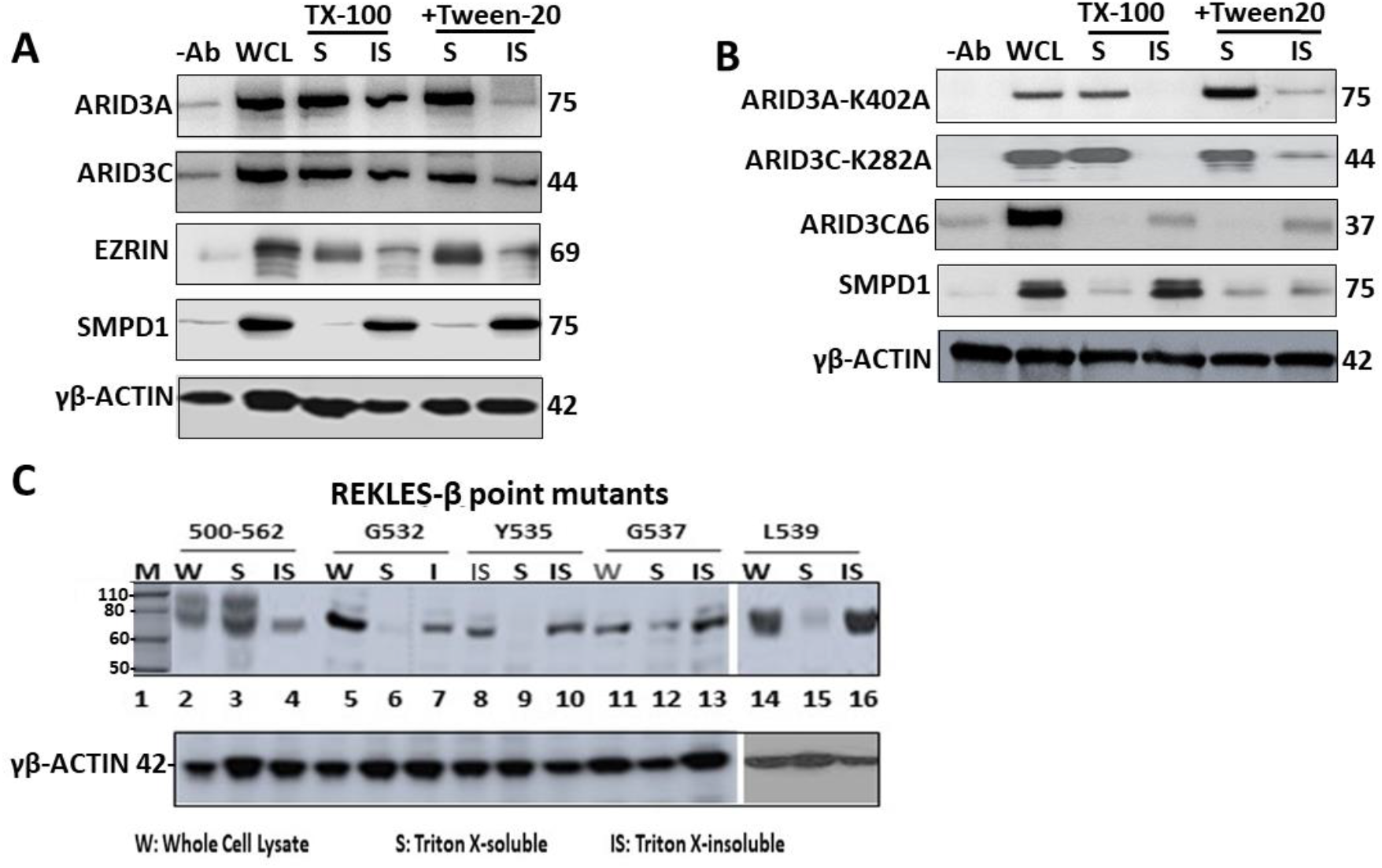
ARID3A and related paralogue, ARID3C localize within monomeric G-Actin via the conserved REKLES-β domain. **A**. ARID3A and 3C transfected COS7 lysates were processed via Triton X-100 and/or TritonX-100 + TWEEN-20 fractionation into insoluble (IS, F-Actin) or soluble (S, GActin) proteins. After fractionation by SDS-PAGE, samples were Western blotted with the indicated Abs; anti-SMPD1 served as an IS positive control and Actin inputs were equalized with pan-Actin Ab. ARID3A and 3C partitioned primarily within the G-Actin fraction. **B**. Loss of the REKLES domain (ARID3A-d(521541) and ARID3CΔ6), but not SUMO-I binding motifs (ARID3A-K402A or ARID3C-K284A) abolishes partitioning into the soluble G-Actin fraction. **C**. A region within REKLES-β (amino acids 532-540 and indicated on S-Figure 5B) contains a cluster of conserved residues (G532, Y535, G537, and L539) whose point mutation significantly reduce or abolish ARID3A association with G-Actin.

### SUMO-modification does not control ARID3 partitioning within G-ACTIN

We previously demonstrated that for BCR signaling, ARID3A requires SUMOylation at a single, conserved lysine residue (K402) for exiting lipid rafts (18; S-Figure 5C). ARID3C requires SUMOylation of the equivalent K284 for rafts exit (18, 29). As shown in Figure 3B, both ARID3A-K402A and ARID3C-K284A fractionate primarily within the soluble fraction. Thus, SUMOylation plays no major role in G-Actin association.

### The REKLES-β domain is essential for ARID3A association within G-Actin

The REKLES domain is named for a hexapeptide (boxed in S-Fig. 5A) that is conserved among all ARID paralogues (30, 31). ARID3C encodes an additional a splice variant (termed ARID3CΔ6) which lacks the C-terminal portion of the highly conserved protein-protein interaction (REKLES-β; exon 6; S-Fig. 6). ARID3CΔ6 fails to undergo nuclear-cytoplasmic shuttling nor does it associate with ARID3A in solution (29). Yet it does bind to common ARID3A DNA binding sites *in vitro* (29). As shown in Figure 3B, loss of this portion of the REKLES domain results in transfer of ARID3CΔ6 to the insoluble fraction.

Sequence alignments allow the REKLES domain to be divided into three sub-domains: A conserved 17 amino acid N-terminal REKLES-α, a relatively conserved 51 amino acid Spacer-region and a highly conserved 59 amino acid C-terminal REKLES-β domain (S-Fig. 5A, B). REKLES-α is required for nuclear localization (NLS) and REKLES-β is essential for nuclear export (NES) (18, 30). REKLES-β also is essential for ARID3A and ARID3C heteromeric interactions (29, 31).

The ARID3CΔ6 observation suggested that the corresponding region of ARID3A, the REKLES-β domain, is critical for ARID3A residence within G-Actin. To test, we first determined if ARID3A carrying a complete deletion (d) (amino acids 500-562; S-Fig. 5C) of the 63 residue REKLES-β was also transferred to the soluble G-Actin fraction. Accordingly, only ∼40% of REKLES β-deficient d500-562 remained within the G-Actin fraction. REKLES-β amino acids 521-541 contain a cluster of residues (G532, Y535, G537, and L539) conserved in all ARID3 paralogues (S-Fig. 5B; 27). Alanine substitutions within each abolished G-protein soluble fraction association to varying degrees.

### The REKLES-β domain is essential for EZRIN-ARID3A interaction

The data above suggested that ARID3A interaction within PM G-Actin is mediated through its REKLES-β subdomain, potentially via interaction with the cytoskeletal linker protein EZRIN. This hypothesis was tested by the Co-Immunoprecipitation (Co-IP) experiments of Figure 4. We transfected COS7 cells with full-length ARID3A (residues 1-601) or with each of the following REKLES mutations: d453-500, in which the entire REKLES domain is deleted (d); d500-521, in which the C-terminal 21 residues of the REKLES-α-REKLES-β Spacer reside just N-terminal to the REKLES-β domain is deleted; t1-541, in which ARID3A is truncated (t) just N-terminal to REKLES-β; and d521-541, in which the REKLES-β domain is deleted (Fig. 4A).

**Figure 4.**
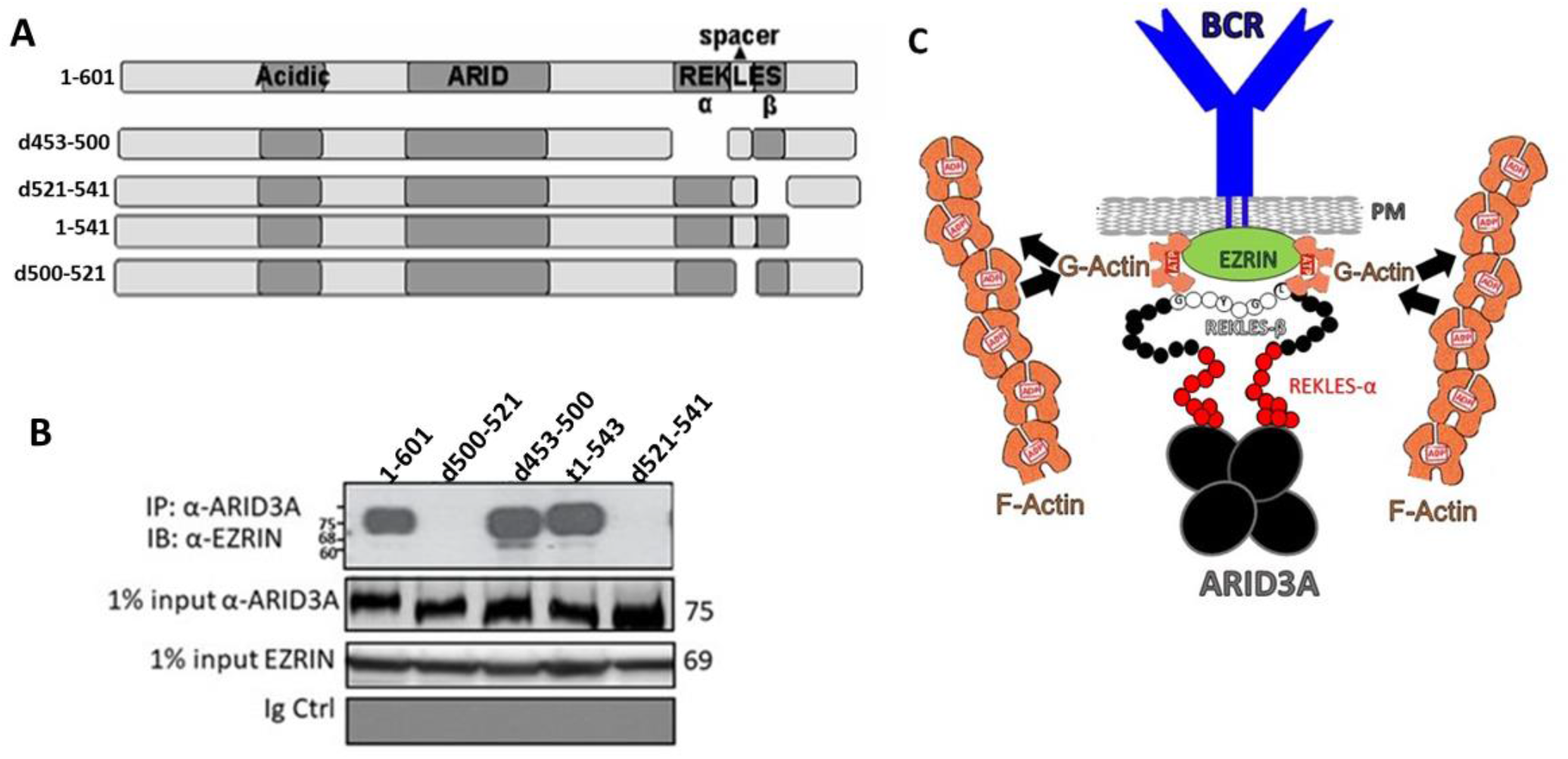
REKLES-β deletions abrogate ARID3A-EZRIN interaction. **A**. Diagram of ARID3A REKLES deletion (d) mutants. Deletions are indicated by gaps. **B**. EZRIN-ARID3A interaction was abrogated only by deletion (d453-500 and d521-541) of the REKLES-β domain. Full length ARID3A and its deletion mutants of (A) were transfected into COS7 cells and lysates were immunoprecipitated with anti-ARID3A prior to resolution on SDS-PAGE followed by anti-EZRIN Western blotting (<u>upper panel</u>); normalization of equivalent inputs (<u>middle panels</u>); anti-EZRIN background control (<u>lower panel</u>). **C**. ARID3A is tethered to G-Actin via interaction of its REKLES-β domain with EZRIN. Cartoon summarizing and extending the findings of this report. We speculate that when the ARID3A tetramer (large black spheres) interacts with EZRIN (green sphere) via a patch of conserved REKLES-β residues (small black spheres with single amino acid codes in white) and just proximal Spacer residues (small red spheres) EZRIN is discharged from lipid rafts in order to release lipid rafts-associated ATP-G-Actin (brown semicircles), possibly for polymerization to ADP-F-Actin (brown chains).

Transfected lysates were immunoprecipitated with a polyclonal anti-ARID3A Ab which previously had been shown to pull-down each of the input mutants (30, 31). After adjusting for equivalent levels of immunoprecipitated ARID3A inputs (Fig. 4B, middle panel), WT and mutant lysates were fractionated on SDS-PAGE and then Western blotted with an EZRIN monoclonal (m) Ab. As shown in Figure 4B, EZRINARID3A interaction was abrogated only by deletion of the REKLES-β domain and N terminal proximal section of the Spacer region.

Collectively these results indicate to us that ARID3A is tethered to G-Actin via interaction of its RECKLES-β and proximal Spacer domain with EZRIN.

## SUMMARY

To our knowledge, ARID3A was the first TF shown to function in lipid rafts (18), but its residence there is not unprecedented. Both STAT1 and STAT3 (33) as well as an isoform of the IgH coactivator OCA B (34, 35) localize to lipid rafts. Several other TFs, including TFII_I, form cytoplasmic complexes (14-17, 36). That TFs function directly or as intermediates in receptor signaling is supported by the association of p35-OcaB with galectin-1 (34) and, in this report, ARID3A with EZRIN.

Previous reports suggested that EZRIN binds specifically to F-Actin, which in turn, increases BCR signaling strength by aggregating and releasing lipid rafts from the cortical Actin cytoskeleton (11, 12). Here we have extended these results by the observation that an ARID3A-EZRIN-Actin interaction occurs exclusively within lipid rafts, and they are co-discharged from rafts following BCR stimulation (Figs. 1, 2, S-Fig. 2). Our data further indicate that, within lipid rafts, EZRIN links ARID3A indirectly to the cytoskeleton to control BCR signaling strength (Figs. 1, 2). But unexpectedly, ARID3A associates almost exclusively with unpolymerized G-Actin (Fig 3A, B). The interaction requires a patch of 5 residues conserved among ARID3 paralogues (S-Fig. 5B) within the 59 residue carboxyl terminal (β subregion) of the multifunctional REKLES domain. In addition, REKLES-β, as well as the C-terminal 21 residues of the REKLES-αβ Spacer, link ARID3A to EZRIN (Fig. 4B).

These results add another layer of complexity to the ARID3A REKLES domain. The REKLES-α subregion encodes an NLS and a binding site for SUMO-I, whereas its β subregion provides essential residues for nuclear export as well as for heteromeric interactions with EZRIN, BTK, PIAS-1 and UBC9 (18, 29-31). The 3-dimensional structure of the DNA binding domain of ARID3A has been solved (37, 38). Yet the structure of REKLES remains to be determined. The observed extent of REKLES multifunctionality as well as the architecture of the platform tethering G-Actin to the REKLES-EZRIN complex is a future goal of our research.

We have combined several of these features into the model of Figure 4C. We suggest that a functional consequence of ARID3A discharge from lipid rafts is to release lipid rafts-associated G-Actin for polymerization to F-Actin. The extent to which this release is achieved contributes to the signaling threshold of the BCR. Conversely, reassembly of the polymerized F-Actin cytoskeleton might act to stabilize interactions among the BCR and additional signaling molecules via trapping the lipid raftlocalized signaling complex. In support, TIRFM studies (capable of observing both the BCR, Actin and EZRIN simultaneously) revealed EZRIN-Actin networks in resting B cells (39). That Ag-Ab binding induces transient EZRIN dephosphorylation followed by detachment of lipid rafts from the Actin cytoskeleton (39) is consistent with our observations (Figs.1, 2) in respect to promotion of BCR-lipid raft interactions. We found it interesting in this regard that ARID3A and EZRIN share a binding partner, S100P. Dimeric S100P binds to and activates EZRIN by unmasking its F-Actin binding sites (40), whereas monomeric S100P represses the DNA-binding and transactivation activity of ARID3A (32).

We feel that further experimental refinement of our model is justified from a health perspective. For example, Diffuse Large B Cell Lymphoma (DLBCL)—both of the <u>A</u>ctivated <u>B C</u>ell (ABC) and <u>G</u>erminal <u>C</u>enter (GC) subtypes—co-segregate with high levels of phospho-EZRIN (41). These results implicate the cytoskeletal network in general, and EZRIN, in particular, in leukemia pathogenesis. Previously we demonstrated that ARID3A is highly expressed in ABC-DLBCL (42). Further, shRNA-mediated ARID3A KD in ABC-DLBCL tumors, but not in GC-DLBCL subsets, led to loss of tumor growth and proliferation (42). While far less is known about ARID3C, RNA-Seq analyses recently identified it within a signaling complex consisting of Protein Tyrosine Phosphatase Receptor type R (PTPRR), α-catenin, β-catenin and E-cadherin that formed exclusively in ovarian cancer (43). These data suggest that modulation of EZRIN, ARID3A, and potentially, ARID3C may provide both prognostic and therapeutic options for these and other malignancies.

## ACKNOWLEGEMENTS

We thank June Harriss, Debora Lerner, Chhaya Das and Maya Ghosh for help in cell culture and molecular techniques. We thank members of the Tucker laboratory for discussions and reading of the manuscript. We received outstanding experimental support at the MD Anderson Smithville Core Facilities, directed by Dr. Jianjun (J-J) Shen, from the following employees: Ms Luis Coletta, Ms Melissa Simper, Dr. Yueping Chen, Ms. Yoko Takata and Dr. Carol Mikulec. H.O.T. received support from NIH Grant R01CA31534, Cancer Prevention Research Institute of Texas (CPRIT) Grants RP100612, RP120348; and the Marie Betzner Morrow Centennial Endowment.

## ARTHUR CONTRIBUTIONS

CS and HOT designed research; CS, LC, JT and DK performed research; CS and HOT analyzed data; HOT wrote the manuscript.

## MATERIALS AND METHODS

### Cell lines

Raji (EBV+, 45); Daudi (EBV+, 45); Ramos (EBV−, 46) and CL01 (EBV−, 47) were obtained from ATCC (Manassas, Virginia) and maintained as previously described (18). Cells were grown and maintained in either Dulbecco’s modified Eagle medium (DMEM) supplemented with 10% fetal bovine serum (FBS; Invitrogen) or in RPMI medium containing 10% FBS. CD43^-^ B cells were prepared by negative selection of whole human blood (Gulf Coast Regional Blood Center, Houston, Texas) or from 10-wk-old BALB/c murine splenocytes (19 30).

### Constructs

Mutant forms of ARID3A and ARID3C used in this study were generated previously by site directed mutagenesis (18, 29-31).

### Preparation of stable sh-RNA retrovirally transduced B cell lines

Stable transductants were established by employment of the Phoenix-A retroviral system. Approximately 3 X10^5^ amphitrophic Phoenix-A packaging cells in 4 ml of DMEM were supplemented with 10% fetal bovine serum (FBS) in 60-mm plates. After one day of culture, cells were transfected using FuGene6, and viral supernatant was harvested 2 days post-transfection, centrifuged, and filtered to remove live cells and debris. The target cell lines described above were plated (3 × 10^5^) onto 60-mm plates and growth medium was replaced with viral mixture. To introduce EZIN-knockdown sequences, we used oligos ordered from Integrated DNA Technologies with restriction site overhangs BbsI and XhoI (sense, 5′-ACCG GCCGTGGAGAGAGAGAAAGATTCAAGAGATCTTTCTCTCTCTCCACGGCTTTTTTACCGGTC-3’; and anti-sense, 5′-TCGAGACCGGTAAAAAAGCCGTGGAGAGAGAGAAAGATCTCTTGAATCTTTCTCTCTCTCCACGGC-3′). Stable lines were established by selection with 2 μg/ml of puromycin from day 2 post-infection.

### B-cell stimulation

To measure signaling effects at low doses of anti-IgM where receptor internalization is minimized, we used monoclonal anti-IgM antibodies in the absence of secondary crosslinking. To stimulate B cells, 500 ng of F(ab’)_2_ fragments of α-μ (clone JDC-15; Dako [α-human]; OB1022; Southern Biotech [α-mouse]) and α-CD19 (clone HD37; Dako [α-human]; clone SJ25-C1 [α-mouse]) were added to 5 × 10^8^ cells for 5 min at 37°. We determined by FACS analysis and semi-quantitative western of lipid raft-associated mIgM (data not shown and S-Figure 1) that under these conditions 1–5% of mIgM in rafts and membranes are engaged.

### Preparation of lipid rafts

Approximately 500 mg of wet cell pellet were washed twice in ice-cold phosphate buffered solution (PBS) and homogenized in 5 ml of 10 mM Tris/Cl (pH 7.4), 1 mM EDTA, 250 mM sucrose, 1 mM phenylmethylsulfonyl fluoride and 1 μg/ml leupeptin (all from Sigma, St Louis, MO) in a tightly fitted Dounce homogenizer using five strokes. The resulting homogenate was centrifuged at 900 *g* for 10 min at 4°C, the resulting supernatant was then subjected to centrifugation at 110 000 *g* for 90 min at 4°C). The resulting membrane pellet was resuspended in ice cold 500 μl TNE buffer (10 mM Tris/Cl [pH 7.4], 150 mM NaCl, 5 mM EDTA, 1% Triton X-100 [Sigma], 10X protease inhibitors [Complete tablets, Roche, Indianapolis, IN]). Sucrose gradients for the preparation of lipid rafts were assembled previously described (18). Lipid rafts were isolated by flotation on discontinuous sucrose gradients. Membrane pellets were extracted for 30 min on ice in TNE buffer. For the discontinuous sucrose gradient, 1 ml of cleared supernatant was mixed with 1 ml of 85% sucrose in TNE and transferred to the bottom of an ultracentrifugation tube, followed by overlay with 6 ml of 35% sucrose in TNE and 3.5 ml of 5% sucrose in TNE. Samples were spun at 200,000 *g* for 30 h at 4°C; fractions were collected from the top of the gradient and analyzed using Western blotting and/or coimmunoprecipitation), as described (30, 31).

### Immunoprecipitation/Western analyses

We employed a stringent RIPA formulation of 500 mM NaCl; 10 mM Tris–Cl pH 8; 0.1% SDS; 5 mM EDTA, pH 8; 10X protease inhibitor (Complete tablet, Roche) to solubilize lipid rafts for subsequent immunoprecipitation experiments. Briefly, buoyant fractions, taken from the discontinuous gradient centrifugation, were pooled and incubated with the same volume of RIPA buffer on ice for 15 min. Resulting extracts were pre-cleared by rocking with 1 ml of a 5% slurry of RIPA equilibrated Protein A beads CL-4B (Amersham Pharmacia, Uppsala) for 4 h at 4°C and removal of the precipitate. The resulting supernatant was then subjected to IP/western assays. The following antibodies were used: α-CD19 (clone 6D5, Dako), α-ARID3A polyclonal Ab (produced in house; 19), α-IgM (BD Pharmingen), α-V5 (Sigma), α-Raftlin (graciously provided by Dr. Akihiko Yoshimura; 21), α-SUMO-1 (Sigma), anti-TFII-I (kindly provided by Dr. Carol Webb,18); pan α-Actin (rabbit origin, Cytoskeleton, Inc), α-Acid sphingomyelinase (SMPD1; ab83354, Abcam), goat α-ezrin (sc-6407; Santa Cruz)

### Accumulation of cytosolic calcium

Indo-1 AM (acetoxymethyl ester; Invitrogen) was added to 3 × 10^6^ leukemia/lymphoma cells in 500 μl Hank’s Balanced Salt Solution (HBSS; Invitrogen) and 10% FCS (HBSS-10). The Indo-1 final concentration was 1 μM. Following incubation at 37°C for 30 min, cells were kept at RT for 5 min and then washed with HBSS containing 2 mM HEPES buffer and no serum (HBSH-0). All manipulations including the incubation were in the dark. Cells were then reacted with a rat anti-human CD16/CD32 Fc blocking Ab (clone 2.4G2, Pharmingen) and washed with HBSH-0 in the dark at 4°C. The cells were resuspended in 50 μl HBSH-0 added to a total volume of 150 μl. Prior to Ca2+ analyses, cells were filtered at RT, warmed to 37°C and then placed in a flow cytometer maintained at 37°C at a flow rate of 250 cells/s. Anti-IgM and anti-CD19 were added (detailed in above in “B Cell stimulation”) 30 s prior to initiation of the experiments. Data acquisition employed a 30-s baseline and was continued for 300 s at 37°C. Cells were analyzed at ∼250 cells/s, and their flux in calcium concentration was determined by 485/405 nm emission ration with excitation at 355 nm. Calibration was performed by measuring Rmin and Rmax, and applying the equation described previously (48). Responses are reported as [Ca2] vs time.

### Purification of Actin

To obtain detergent-soluble (Actin-free) and insoluble (Actin-enriched) fractions, ARID3A-transfected COS7 cells were harvested by scraping, and then resuspended in 750 μl of pre-warmed LAS buffer [50 mM Pipes (pH 6.9), 50 mM NaCl, 5 mM MgCl2, 5 mM EGTA, 5% (v/v) glycerol, 0.1% NP-40, 0.1% Triton X-100 (with or without 0.1% Tween 20), 0.1% β-mercaptoethanol, 1 mM ATP, 0.001% Antifoam C, and a protease inhibitor cocktail consisting of 0.4 mM tosyl arginine methyl ester, 1.5 mM leupeptin, 1 mM pepstatin A, and 1 mM benzamidine]. Cells were lysed by 10 passages through a 25-gauge needle. The lysate was clarified by centrifugation (400*g* for 5 min at RT). The supernatant (100 μl) was collected, and Actin polymerization was initiated by addition of 2 mM MgCl2, 0.2 mM EGTA, 100 mM KCl, and 1 μM phalloidin and by incubation for 1 hour at 37°C. F-Actin and F-Actin– binding proteins were pelleted by ultracentrifugation (270,000*g*) for 1 hour at 4°C. The supernatant (GActin fraction) was collected, and the pellet (F-Actin fraction) was washed twice with Milli-Q water and resuspended in 100 μl of 8 M urea. After addition of SDS loading buffer, the samples were separated by SDS–polyacrylamide gel electrophoresis (PAGE) and analyzed by Western blotting for Actin using anti– pan-Actin polyclonal rabbit Ab (1:2000; Cell Signaling Technology, 4968).

### Immunofluorescence staining of murine B cells

Mouse CB7Bl/6 MZ and FO B cells were isolated from mouse spleen cell suspension by anti-CD19 exclusion using a kit #130-100-366 (Miltenyi Biotec). Cells were fluorescently stained with rabbit αβ-Actin mAb (SP124; ab115777; Abcam) developed with goat anti-rabbit IgG H&L (Alexa Fluor® 594), mouse anti-ARID3A mAb A-4 (sc-398367, Santa Cruz Biotechnology) developed with goat anti-mouse IgG H&L (Alexa Fluor® 488; ab150117, Abcam) and stained with DAPI (Staining Solution ab228549, Abcam). Visualization is described below and shown in Figure 2B and S-Figure 4.

### Multiphoton microscopy (MPM) imaging

MPM imaging as detailed by Kuhn and Poenie (48) was performed at a rate of one to two processed frames per second. Each image in the resulting MPM movie sequence was enhanced slightly using a 3 × 3 convolution kernel of [[− 1/2 1/2 1/2], [− 1 1/2 1/2], [− 1/2 1/2 1/2]], which added a small emboss effect to the image and improved visibility. Fluorescent images were acquired using a 12 bit CCD camera (Model DVC-1312M, DVC, Austin, TX) on a Nikon Diaphot 200 fluorescence microscope. Image stacks were obtained using a MAC 2000 z-axis focus controller (Ludl Electronic Products, Hawthorne, NY) and a custom image acquisition plugin written for ImageJ. The point-spread function (PSF) was measured under similar conditions using 100 nm fluorescent beads (L-5473, Molecular Probes) diluted in dH2O and dried on glass slides to give approximately one bead per field. Image stacks were deconvolved for 500 iterations using the expectation maximization algorithm in XCOSM (49). 3D projections were calculated using the maximum value projection method in ImageJ, and stereo pairs were generated using images that differed in rotation by 10 degrees. Microtubule images were enhanced using a multiscale 3D line filter implemented as a custom plugin for ImageJ.

## SUPPLEMENTAL FIGURE LEGENDS

**S-Figure 1.**
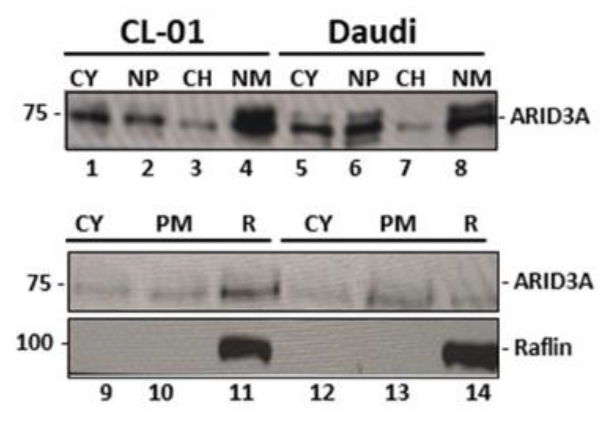
Biochemical fractionation indicates that lipid rafts, but not other subcellular compartments, vary among B cell tumors in levels of ARID3A. Exponentially growing Daudi and CL01 cells were fractionated into cytoplasm (CY), soluble nucleoplasm (NP), chromatin (CH) and nuclear matrix (NM) according to previous protocols (18, 27). Immunoblotting of equal aliquots (∼25 µg) of protein with anti-ARID3A antiserum (Lanes 1-8) revealed no differences between these two cell lines. Cytoplasmic extracts (Lanes 1, 9, 5 and 12; 25µg as equivalent to ∼2 × 10^5^ cells) were used to normalize Western (Lanes 9-14,). Plasma membranes (PM; lanes 10 and 13) and lipid rafts (R; lanes 11 and 14) were prepared from ∼1 × 10^7^ cells, as described above. The entire preparation was used for SDSPAGE/Western, which was internally normalized by reprobing the filter with anti-Raflin antiserum.

**S-Figure 2.**
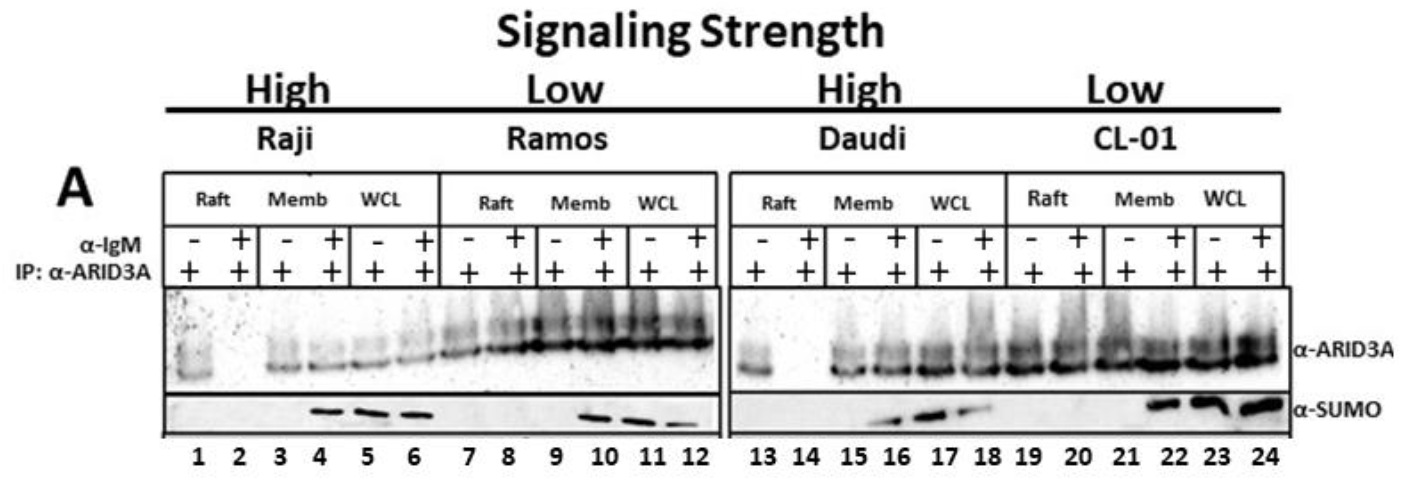
Strength of BCR stimulation leads to differential lipid rafts discharge of ARID3A, but not its interacting partner, SUMO-I. SUMO-I levels were assessed under conditions in which lipid rafts levels of ARID3A were modulated as described in the text and in Figure 1. **A**. Following moderate stimulation with anti-IgM, ARID3A is discharged from lipid rafts of Raji and Daudi cells (<u>lanes 2 and 14</u>), but not from those of Ramos and CL01 cells (<u>lanes 8 and 20</u>). **B**. ARID3A remains localized within rafts of all cell lines following weak stimulation by anti-CD19 (<u>lanes 2, 8, 14 and 20</u>). **C**. Under the strongest stimulatory condition (anti-IgM + anti-CD19), ARID3A is fully discharged (<u>lanes 2, 8, 14 and 20</u>). **A-C**. No change was observed in the expression levels of SUMO-1, which modifies ARID3A and ARID3C at single K residues (S-Figure 5C) and interacts with ARID3A and ARID3C in lipid rafts (18)

**S-Figure 3.**
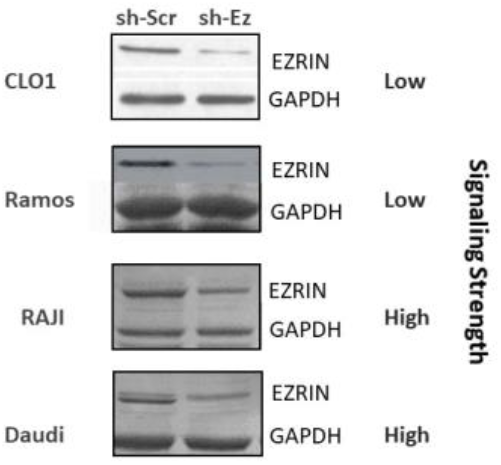
Ezrin retroviral knockdown levels. Levels of ezrin were determined following shEzrin and sh-Scrambled transduced lymphoma/leukemia cells via Western blotting as detailed in Methods and Materials. Data shown are representative of three independent experiments

**S-Figure 4.**
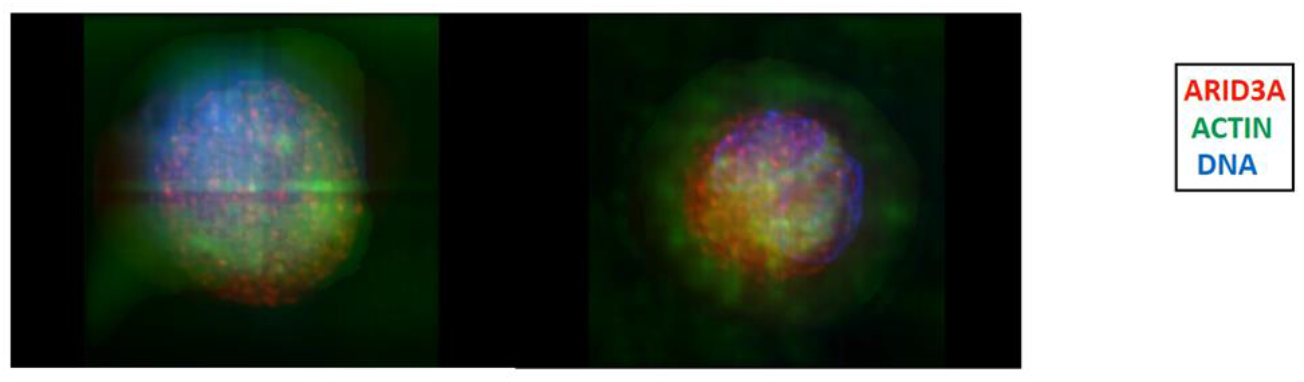
Computerized 3D reconstruction demonstrate Actin-ARID3A interaction in B lymphocytes. Murine FO B cells were purified, stained with fluorescent-antibodies and imaged as detailed in Materials and Methods. Significant colocalisation (yellow) was observed for ARID3A (red) and Actin (green). DAPE-stained nuclei are shown in blue.

**S-Figure 5.**
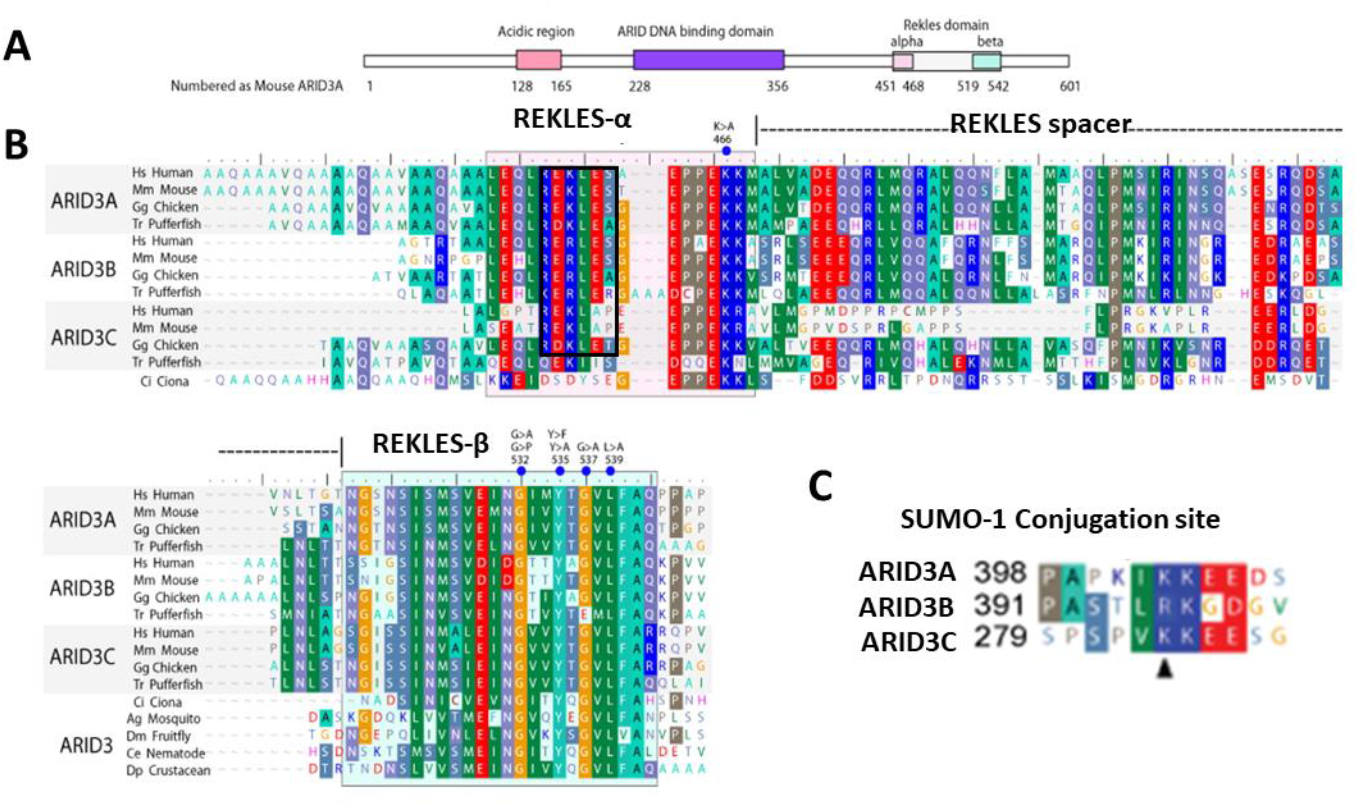
ARID3 family REKLES domain alignments. **A**. Schematic of ARID3A with domains denoted in colored boxes, including the Acidic, DNA binding (ARID) and multifunctional homomerization/nuclear export/EZRIN-interacting REKLES domain. **B**. The alignment of ARID3 REKLES-α subdomains of human (*Homo sapiens*), mouse (*Mus musculus*), chicken (*Gallus gallus*), pufferfish (*Takifugu rubripes*), and the urochordate tunicate Ciona (*Ciona intestinalis*) are shown. For the REKLES-β alignment, additional sequences from mosquito (*Anopheles gambiae*), fruit fly (*Drosophila melanogaster*), nematode (*Caenorhabditis elegans*), and the crustacean Daphnia (*Daphnia pulex*) are indicated. Point mutants used in this study are denoted with a blue dot above the aligned sequence. **C**. A SUMO-I consensus motif with conjugated K (▴) is located N-terminal to the REKLES domain at the indicated positions for ARID3A and ARID3C. Modified from references 29-31.

**S-Figure 6.**
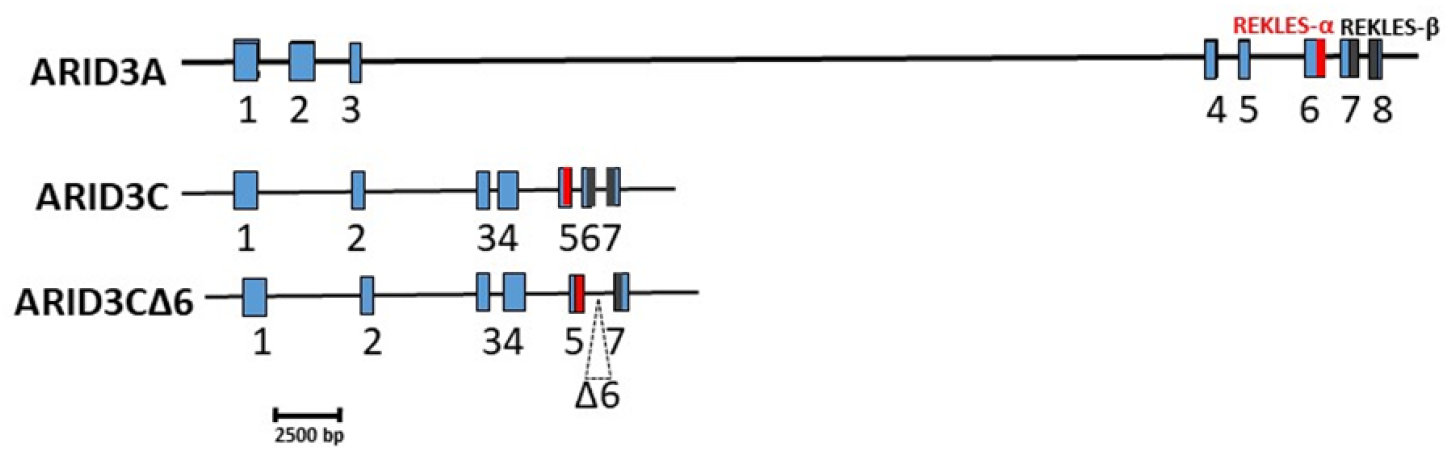
Schematic of ARID3A and ARID3C loci. Shown in scale (kb marker at bottom) are exons encoding ARID3A, ARID3C, and ARID3CΔ6. ARID 3C and ARID3CΔ6 loci are compressed relatively to that of ARID3A. REKLES-α exons, red; REKLES-β exons, gray. The dotted triangle indicates the alternative pre-mRNA skipping of exon 6.

